# Implicit Versus Explicit Timing – Separate or Shared Mechanisms?

**DOI:** 10.1101/2022.03.21.485175

**Authors:** Sophie K Herbst, Jonas Obleser, Virginie van Wassenhove

## Abstract

Time implicitly shapes cognition, but time is also explicitly represented, for instance in the form of durations. Parsimoniously, the brain could use the same mechanisms for implicit and explicit timing. Yet, the evidence has been equivocal, revealing both joint versus separate signatures of timing. Here, we directly compared implicit and explicit timing using magnetoencephalography, whose temporal resolution allows investigating the different stages of the timing processes. Implicit temporal predictability was induced in an auditory paradigm by a manipulation of the foreperiod. Participants received two consecutive task instructions: discriminate pitch (indirect measure of implicit timing) or duration (direct measure of explicit timing). The results show that the human brain efficiently extracts implicit temporal statistics of sensory environments, to enhance the behavioral and neural responses to auditory stimuli, but that those temporal predictions did not improve explicit timing. In both tasks, attentional orienting in time during predictive foreperiods was indexed by an increase in alpha power over visual and parietal areas. Furthermore, pre-target induced beta power in sensorimotor and parietal areas increased during implicit compared to explicit timing, in line with the suggested role for beta oscillations in temporal prediction. Interestingly, no distinct neural dynamics emerged when participants explicitly paid attention to time, compared to implicit timing. Our work thus indicates that implicit timing shapes the behavioral and sensory response in an automatic way, and is reflected in oscillatory neural dynamics, while the translation of implicit temporal statistics to explicit durations remains somewhat inconclusive, possibly due to the more abstract nature of this task.

## Introduction

Implicit and explicit timing orchestrate perception and action, allowing us to interact with dynamic environments (Coull et al., 2011, 2013; Coull and Nobre, 2008; Michon, 1990). Dynamic sensory environments have a temporal structure, defined for example by delays between events, order, or synchrony. Such temporal features can be learned to form temporal predictions about future events, a mainly implicit or unconscious cognitive process. At the same time, temporal features can be perceived explicitly and consciously, for example time dragging when waiting for the bus.

In accordance with the literature, we adhere to the following working definitions: *Implicit timing* is defined as the extraction of temporal contingencies between perceived events, resulting in facilitation of behaviour (indirect measurement). *Explicit timing* is defined as the deliberate engagement in timing, resulting in overt temporal estimates (direct measurement).

It is currently an open question whether the brain uses the same mechanisms for the online coding of elapsed time for implicit and explicit timing, approached mainly as separate processes in previous studies. Behavioural studies provided some evidence for shared mechanisms: both implicit and explicit timing are subject to scalar variability (Piras and Coull, 2011; but see Ameqrane et al., 2014), and behavioural measures of implicit and explicit timing partially correlate over participants (Coull et al., 2013, but only for long intervals). However, several recent studies show dissociable patterns of responses in implicit versus explicit timing tasks (Droit-Volet and Coull, 2016; Droit-Volet et al., 2019; Los and Horoufchin, 2011; Mioni et al., 2018). In sum, these results suggest that timing mechanisms are at least partially task-specific, but it is currently unclear which cognitive and neural processes are responsible for the divergence.

Considering the anatomical networks underlying implicit and explicit timing, an influential set of studies directly compared both tasks using fMRI (Coull et al., 2013; Coull and Nobre, 2008), and found a common substrate in pre-motor areas, but distinct contributions to implicit timing from the left inferior parietal cortex and the cerebellum, versus, for explicit timing, from the right frontal and parietal areas, and the basal ganglia. The anatomical sources identified by studies performed separately on implicit timing (using fMRI or EEG/ MEG; Herbst et al., 2018; Martin et al., 2006; Mento et al., 2013; Praamstra et al., 2006) partially converge with those well known for explicit timing, namely pre-motor and motor areas (Wiener et al., 2010). However, implicit timing also engages brain networks more generally associated with attention and predictive processing in frontal and parietal cortices (Coull and Nobre, 2008; Meindertsma et al., 2018; Visalli et al., 2019), as well as the cerebellum (Breska and Ivry, 2018).

A large number of EEG and MEG studies have investigated the neural dynamics of implicit and explicit timing, implicating all canonical frequency bands (see review by Wiener and Kanai, 2016). The strongest overlap between studies on implicit and explicit timing occurs in the modulation of intermediate to high frequent oscillatory power (alpha/ 8–12 Hz, beta/ 15–30 Hz, gamma/ 30–100 Hz), described in both, implicit (Cravo et al., 2011; Meindertsma et al., 2018; Todorovic et al., 2015), and explicit (Kononowicz and Rijn, 2015; Kononowicz et al., 2018; Kulashekhar et al., 2016; Spitzer et al., 2014; Wiener et al., 2018) timing situations. In particular, beta power fluctuations have been observed over a range of different paradigms, but their distinct role for unique aspects of temporal cognition remains to be established.

Here, we set out to directly compare implicit and explicit timing using magnetoencephalography (MEG), based on the same sensory inputs. Participants were asked to perform either a pitch discrimination task or a duration discrimination task during an auditory foreperiod paradigm (Niemi and Näätänen, 1981; adapted from Herbst and Obleser, 2019). To induce implicit timing, we manipulated the foreperiod over blocks, such that it was either predictive (same foreperiod throughout the block) or non-predictive (variable foreperiod throughout the block). Combining the manipulation of predictability with the two tasks resulted in a balanced 2 × 2 factorial design (see Fig.1A, B), with the factors **task** (pitch versus duration discrimination) and **predictability** (predictive versus non-predictive), to address how the same temporal statistics inform implicit predictions versus explicit temporal decision making.

**Figure 1:**
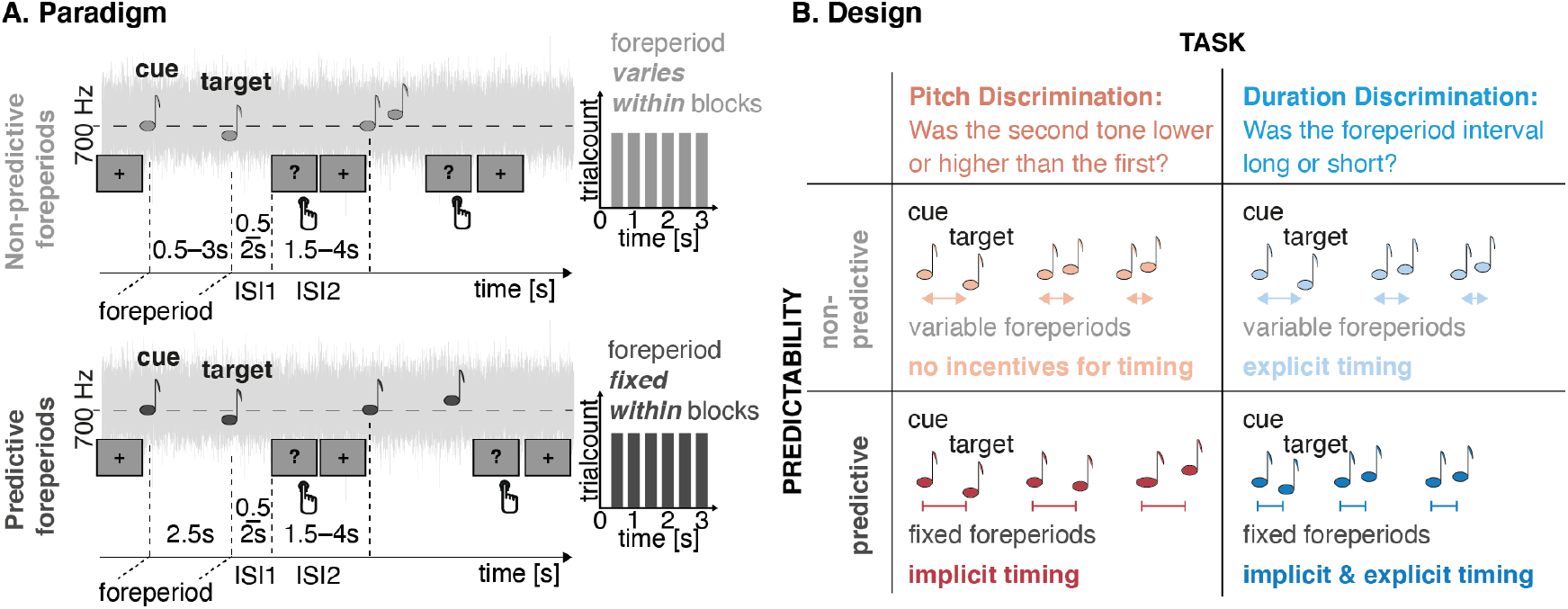
Experimental design: **A. Foreperiod paradigm:** tone pairs (cue and target) were presented, embedded in white noise. Unbeknownst to participants, temporal predictability was manipulated block-wise, by either varying the foreperiod within a block (non-predictive condition, light grey, upper panel), or using a fixed foreperiod throughout a block (predictive condition, dark grey, lower panel). Overall, the same foreperiods were used in both conditions. **B. Design:** in a 2 x 2 design, the predictability manipulation was crossed with two tasks: participants were either asked to judge whether the pitch of the target tone was higher or lower than the one of the cue (pitch discrimination task), or whether the foreperiod interval was short or long (duration discrimination task), in comparison to all previously presented intervals.

Importantly, this design allowed us to compare implicit and explicit timing during the foreperiod: in the pitch discrimination task, participants could anticipate the onset of the target, hence implicitly time the foreperiod. In the explicit timing task, participants could also anticipate the target based on the same temporal statistics, but, additionally, had to explicitly encode the duration of the foreperiod. Our results show that participants systematically formed temporal predictions from the temporal statistics of the inputs, resulting in enhanced behavioral and neural processing of the auditory stimuli. Orienting attention in time was reflected by an anticipatory increase of alpha power over parietal and visual areas in the temporally predictive conditions in both tasks. Beta power was increased during implicit compared to explicit timing during the foreperiod, in line with the previously proposed role of beta oscillations for temporal prediction.

## Methods

### Participants

Twenty-four right-handed volunteers (mean age: 24.4, SD: 4.4 years, 11 females) with no self-reported hearing loss or neurological disorder participated in the experiment. The sample size was planned based on a previous EEG study using a very similar foreperiod paradigm (N = 24, Herbst and Obleser, 2019). Each participant gave written informed consent prior to the experiment in accordance with the Declaration of Helsinki and the Ethics Committee on Human Research at Neurospin (Gif-sur-Yvette). The data of one participant was completely removed, due to technical problems resulting in missing blocks. The MEG data of four participants was excluded from the MEG analysis due to technical problems with the head positioning system during MEG-acquisition. The behavioral data of these four participants were kept in the analysis.

### Software and Toolboxes

The experimental protocol was implemented in the *Psychophysics Toolbox* (Version 3.0.13 Brainard, 1997; Pelli, 1997) for Matlab (R2017a), run on a laptop under Windows 7. Behavioral analyses were carried out in *Matlab* (version R2019b), using the *Psignifit toolbox* (Version 4, Schütt et al., 2016), as well as in *R* (Team, 2016), using the and the *lme4* package (Bates et al., 2015). MEG analyses were carried out in *MNE python* (Versions 20.0 – 23.0 Gramfort et al., 2013, 2014), in combination with *Freesurfer* for MRI surface reconstruction (Dale et al., 1999).

### Data Availability

Data will be shared upon request.

### Procedure

Participants were recruited at NeuroSpin (Gif sur Yvette, France), for an experimental session of about 4 h, which included an MRI safety screening, the preparation for EEG and MEG recording, 1.5 h of simultaneous M/EEG recording, debriefing, and a subsequent anatomical MRI exam. The M/EEG recording started with short tests to adjust the auditory stimuli (described in more details below). During the M/EEG recording, participants were presented with an auditory foreperiod paradigm (adapted from Herbst and Obleser, 2019), and were asked to perform a pitch discrimination task in half of the session, and a duration discrimination task in the other half. Each participant performed 12 blocks of both tasks in succession with the task order being counterbalanced across participants. Between the two tasks, participants took a short break, but remained seated in the MEG chair. After the full MEG recording, participants received a post-experimental questionnaire, were debriefed, and underwent an anatomical MRI exam, which lasted about 20 minutes.

### Stimuli

Auditory stimuli were delivered via in-ear headphones (EatyMotic). Lowpass (5 kHz) filtered white noise was presented constantly throughout each block, at 50 dB above the individual sensation level, which was determined for the noise alone at the beginning of the experiment using the method of limits. Pure tones of varying frequencies (duration 50 ms with a 10 ms on- and offset ramp) were presented with a tone-to-noise ratio fixed at −18 dB relative to the individual noise level.

The first tone, to which we will refer as the *cue* had a fixed frequency of 700 Hz. The second tone, the *target*, was varied in individually predetermined steps, logarithmically transformed and multiplied with the cue tone’s frequency, to obtain a log-spaced frequency scale around the cue. The frequency steps were predetermined during an initial training phase: each participant was first presented with one short block (18 trials) of the pitch discrimination task, with feedback after each trial, to familiarize themselves with the task. In a second training block, performance was computed over all trials (30), and the training was terminated if the participant reached between 55 – 75% accuracy. If performance was below 55% (too close to chance level) or above 75% (suggesting a possible ceiling effect), the frequency-steps were increased or decreased, respectively, and the participant performed another block of the training. This way, participants underwent minimally two blocks (2.67 blocks on average), and the maximal number of training blocks needed was five (one participant). The frequency-steps were kept as determined by this procedure throughout the whole experiment for both tasks. The foreperiods used in the duration discrimination task were not adjusted individually, because varying intervals would have been difficult to use in the group analyses of the EEG data.

To induce implicit timing, the interval between cue and target tones, *the foreperiod*, was varied in six linear steps from 0.5 s to 3.0 s (see also Fig. 1A). Per experimental block (30 trials), foreperiods were either variable (*temporally non-predictive*), that is each foreperiod interval occurred five times per block in random order, or fixed (*temporally predictive*), that is during one block only one foreperiod interval occurred. In the predictive condition, the fixed foreperiods were varied between blocks, such that one block was presented per task and foreperiod interval. Blocks with fixed and variable foreperiods were presented randomly interleaved. During the training blocks, only variable foreperiod blocks were presented.

After the target tone, participants had to withhold their response for a delay, varying between 0.5 – 2.5 s (exponential distribution, truncated at 2.5 s), until a black question mark was presented on the screen as response prompt. The question mark disappeared when a response was given, or after 2 s, followed by a variable inter-trial interval between 3 – 4.5 s (exponential distribution, truncated at 4.5 s). To respond, participants pressed one out of two coloured buttons on a Fiber Optic Response Pad (FORP, Science Plus Group, DE), using the index or middle finger of their right hand.

### Tasks

Each participant performed both tasks, pitch discrimination and duration discrimination, based on the exact same auditory stimuli. In the pitch discrimination task, participants were instructed to indicate whether the target tone was lower or higher in pitch than the cue tone. In the duration discrimination task, participants had to indicate whether the interval between the cue and target tone was short or long, in comparison to all the intervals presented in the study. They were instructed not to count seconds (Rattat and Droit-Volet, 2012). We did not present a standard interval on each task to keep the stimulation equal to the pitch discrimination task. To make sure that participants were sufficiently exposed to all possible time intervals before starting the main task, we presented variable foreperiod blocks during training (2 – 5 blocks per participant). In the predictive blocks of the duration discrimination task, participants were effectively asked to judge the duration of the same foreperiod interval on every trial. However, no feedback was given per trial, so duration judgements remained subjective. We verified in the behavioral analyses that participants did engage in explicit timing, both in the predictive and non-predictive conditions.

In addition to the training at the beginning of the experimental session, participants performed one short block of training at the start of each task (pitch or duration discrimination, 18 trials). Feedback was provided after each response, by the color of the fixation cross (correct answer: green, wrong answer: red), or by the phrase ‘pas de réponse’ (no response) if the participant did not respond. During the experimental blocks, participants did not receive trial-wise feedback, but were informed after each block about their average performance. The no-response feedback was shown after each missed trial.

Per task, participants performed 12 blocks of 30 trials. Six blocks had random foreperiods (six foreperiods appearing randomly, five times each), and six blocks had fixed foreperiods (one out of the six foreperiods presented constantly throughout the block), resulting in 720 trials overall (excluding tone frequency adjustment and training blocks).

### Post-experimental questionnaire

After the M/EEG recording, participants were asked a set of questions (see Appendix), to rate the subjective difficulty of the tasks, and check whether they had noticed the manipulation of predictability. To achieve the latter, we first asked participants whether they realized that the time intervals between cues and targets varied, and second, whether they could describe the variability. Finally, the experimenter explained the predictability manipulation, and asked whether the participant had noticed it.

### M/EEG recordings

Prior to the arrival of the participant, an empty room recording was performed for 1 minute to assess the noise level of the MEG sensors.

Before the M/EEG recording, participants were equipped with an EEG cap (64 Ag-AgCl electrodes, EasyCap, Germany). The impedances of the electrodes were adjusted below 15 kΩ. External electrodes were positioned to record the electro-occulograms (EOG, horizontal and vertical eye movements) and electro-cardiograms (ECG). The positions of the EEG electrodes, four head-position coils, and three fiducial points (nasion, left and right pre-auricular areas) were digitized using a 3D digitizer (Polhemus, US/Canada) for subsequent co-registration with the individual’s anatomical MRI.

The combined M/EEG recordings took place in a magnetically shielded chamber, where the participant was seated in an arm-chair under the MEG helmet. The electromagnetic brain activity was recorded using a whole-head Elekta Neuromag Vector View 306 MEG system (Neuromag Elekta LTD, Helsinki) with 102 triple-sensors elements (two orthogonal planar gradiometers, and one magnetometer per sensor location) and concurrent recording of electric brain activity (EEG).

Participants were instructed to fixate their gaze on a screen positioned in front of them, at about one meter distance. The chamber was dimly lit. Their head position was measured before each recording run (8 in total) using four head-position coils. M/EEG recordings were sampled online at 1 kHz, and low-pass filtered at 330 Hz, but no online high-pass filter was used.

### Anatomical MRI recordings

Anatomical Magnetic Resonance Imaging (aMRI) was used to provide high-resolution structural images of each individual’s brain. The anatomical MRI was recorded directly after the MEG recording using a Siemens 3 T Magnetom Prisma Fit MRI scanner. Parameters of the sequence were: slice thickness: 1 mm, repetition time TR = 2300 ms, echo time TE = 2.98 ms, and flip angle 9 degrees.

### Analyses

#### Behavioural data

We analysed accuracy as the proportion of correct responses, and response times from the onset of the response prompt to the registered button press (Figure 2A, B). We compared the average accuracy and response times per participant between tasks and predictive versus non-predictive conditions, using paired t-tests. For the response times, we additionally computed a repeated measures ANOVA with the factors foreperiod and predictability. Response times were log-transformed before being entered into the statistical tests.

**Figure 2:**
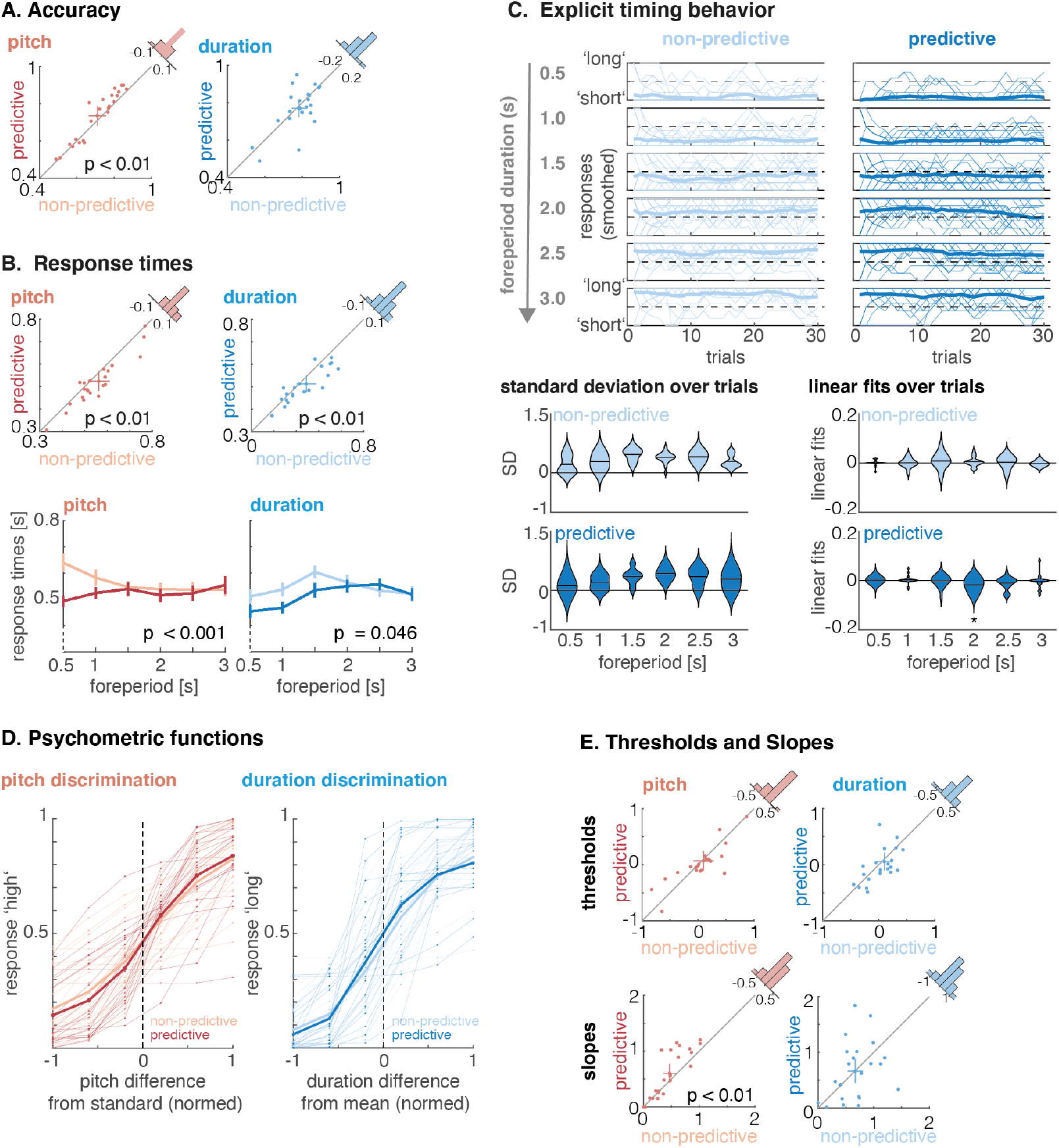
Behavioral results: **A. Accuracy:** temporal predictability increased pitch discrimination accuracy (red scatter plot), but not duration discrimination accuracy (blue scatter plot). The cross depicts the mean and 95% confidence intervals in the directions of both variables. **B. Response times:** response times were faster in the predictive condition in both tasks. Furthermore, we found an interaction between foreperiod and predictability in both tasks: response times of the predictive and non-predictive conditions diverged predominantly at the short foreperiods. Error bars depict the standard error of the mean. **C. Explicit timing behavior:** Top: responses (coded as ‘short’ = −1, ‘long’ = 1, moving average over 5 trials) per condition and foreperiod. Thick lines are the participant average, thin lines depict single participants’ responses. Bottom left: Participants’ standard deviations of responses over trials, per condition and foreperiod. The black horizontal lines depict the average. Bottom right: Linear fits over responses throughout a block. A significant negative slope was found for the 2 s foreperiods in the predictive blocks). **D. Psychometric curves:** pitch discrimination task: the proportion of responding ‘high’ is fitted as a function of the pitch difference between cue and target tone; duration discrimination: the proportion of responding ‘long’ is fitted as a function of the foreperiod duration. The thin lines show the curves of single participants, and the thick line the average thereof. **E. Thresholds and slopes of the psychometric function:** thresholds (upper row) were not affected by the predictability manipulation. In the pitch discrimination task only, the slopes (lower row) were steeper in the temporally predictive condition. No slope difference was found in the duration discrimination task. The cross depicts the mean and 95% confidence intervals in the directions of both variables.

To verify whether participants had engaged in explicit timing, especially also in the predictive blocks when the same interval was presented on each trial, we analysed the variability of participants’ duration discrimination responses over trials per condition (predictive, non-predictive) and foreperiod (Figure 2C). To this end, we first computed a moving average of 5 trials of the responses coded as −1 for ‘short’ and 1 for ‘long’ over the trials per block. Note that the trials combined to one ‘block’ (same foreperiod, same condition) were presented consecutively in the predictive conditions, but taken from different experimental blocks in the non-predictive condition. We then computed participants’ standard deviation per block as a measure of response variability, and assessed linear fits over trials to capture a shift in the reference throughout a block.

To obtain a more refined measure of pitch and duration discrimination sensitivity, a cumulative normal function was fitted to the responses using the *Psignifit toolbox* for Matlab (Version 4, Schütt et al., 2016). We used Psignifit’s default priors for the threshold, slope, guess, and lapse-rates, based on the given stimulus range (Schütt et al., 2016, p.109). To obtain the same stimulus levels for all participants for fitting the psychometric functions, the individually adjusted pitch steps were normed to six linearly spaced steps from −1 to 1, with −1 and 1 reflecting each participant’s lowest/ highest tones, and 0 being the pitch of the standard tone. For the duration discrimination task, foreperiods were also normed between −1 and 1. We fitted psychometric functions to each individual’s data separately per condition and compared the resulting parameters between conditions (threshold, slope, guess- and lapse rates) using two-sided t-tests. Additionally, we calculated Bayes Factors for all statistical tests, using the *Bayes Factors* package for Matlab (Rouder et al., 2009). All behavioral results are depicted in Figure 2D – E.

#### M/EEG preprocessing

M/EEG data were analysed using MNE python, following a standardized pre-processing pipeline ^1^. Due to substantial artefacts, we chose to disregard the EEG data. The following description refers only to the MEG data. As a first step, noisy or flat sensors were identified visually and marked (5.1 on average). The raw MEG data were then pre-processed with Maxfilter implemented in MNE python (using spatiotemporal signal source separation, duration: 30 s, tSSS; Taulu and Simola, 2006), which also interpolates the bad sensors. Head coordinates were read from the 5th of the eight recording runs, and all other runs were spatially aligned to these coordinates.

The continuous data were low-pass filtered at 120 Hz using a finite impulse response filter (with a hamming window and a transition band-width of 30 Hz), run forward and backward. Data were downsampled to 256 Hz, epoched with respect to the cue tones (−3 – 7 s), and linearly detrended over the full 10 s time window.

A custom-build rejection procedure was applied to remove noisy epochs, rejecting epochs that exceeded each participant’s average global field power (standard deviation over either magneto- or gradiometers) by more than three standard deviations. Independent component analysis (ICA) was performed on the remaining epochs, with an additional 1 Hz high-pass filter applied only for that purpose. For the detection of artefacts related to the electroocculo- (EOG) and electrocardiogram (ECG), we used an inbuilt routine in MNE python, which finds a participant’s typical EOG and ECG activity, and returns the ICA components that correlate with these typical events. Thresholds for returning the ICA components were set to: 0.75 for the ECG (Dammers et al., 2008, cross-trial phase statistics), and 3.0 for the EOG (z-score). In addition, ICA components were screened manually and removed if they could be identified as noise (27.57 manually identified components on average).

To remove remaining artefacts, a fixed threshold rejection procedure was applied, removing epochs with amplitudes exceeding 6000fT (magnetometers) and 4000fT/cm (gradiometers). On average, 696.84 epochs (out of 720, SD = 13.79, range = 654 – 710 epochs) were retained per participant. For the analyses of the time window around the target stimulus, we re-aligned epochs to the target-tones (−3.5 – 3 s). We also produced a specific version of the target-locked epochs with the aim to look for a slow ramping activity during the foreperiod. These epochs were low-pass filtered at 7 Hz (a-causal FIR filter, hamming window, applied forward and backward, transition bandwidth 2 Hz) and baseline corrected to −0.5 – 0 s from cue onset.

In the following, we report only the analyses of the 102 magnetometers, to reduce the dimensionality of the data and simplify the interpretation of the results and topographies.

#### Evoked fields

To assess whether task and temporal predictability affect the processing of cue and target stimuli, we analysed the evoked fields in response to the presentation of the tones. Statistical results are depicted in Figure 3 (with an additional 7 Hz low-pass filter applied only for display purposes) and reported in Table 1. The statistical analysis was implemented on two levels. First, we computed a linear regression on the single trial data per participant. Then, we submitted the individual regression weights to spatio-temporal cluster-based permutation tests to test for significance at the group level.

**Figure 3:**
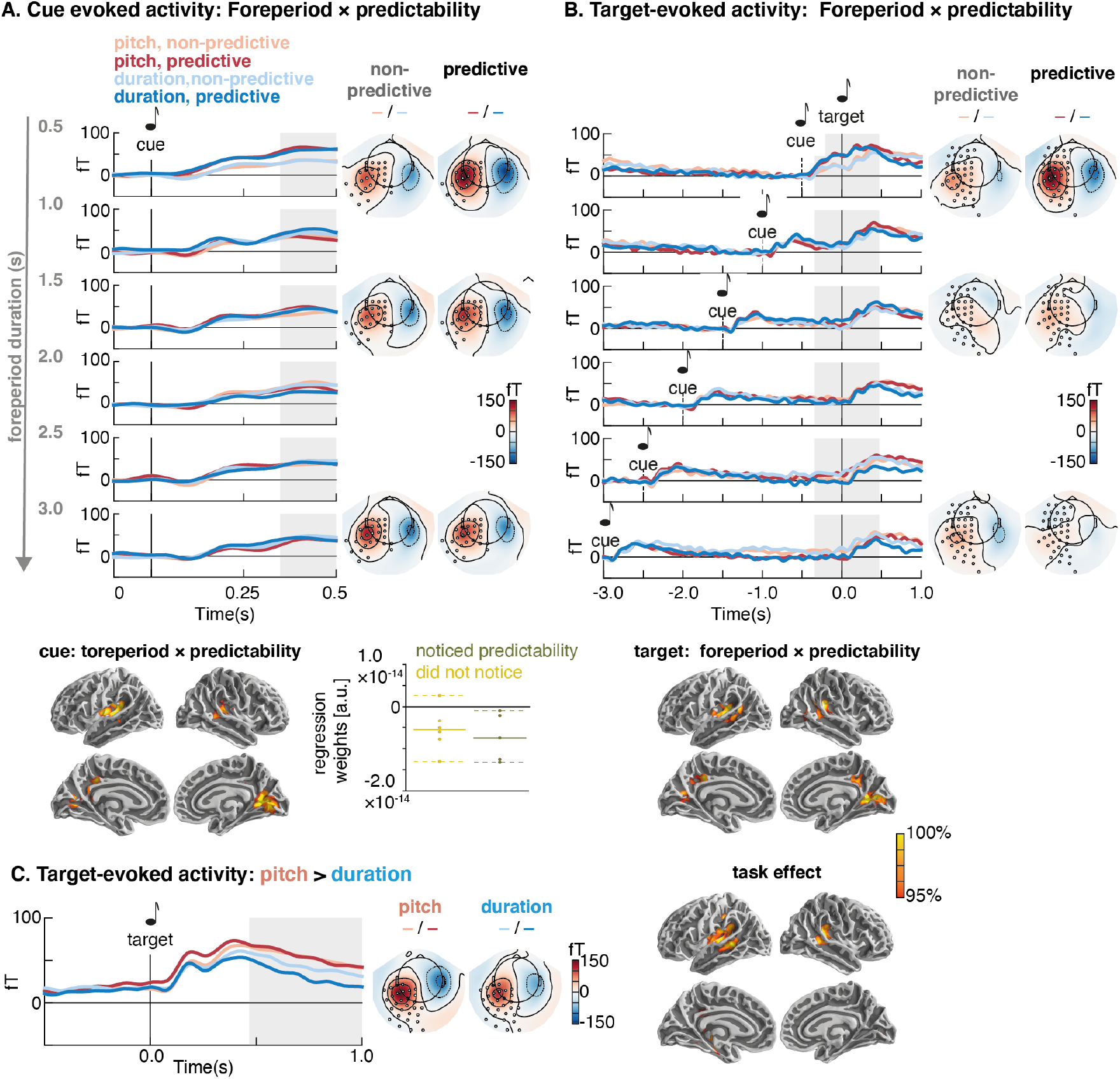
Evoked fields and source estimates: **A. Cue-evoked activity** differed at latencies following temporally predictive vs. non-predictive cues in a post-cue time window of 0.35 – 0.50 s, regardless of the task. The effect was stronger for shorter foreperiods. The sensor-space topographies show the evoked activity, extracted at the clusters that formed the significant cluster (marked by black circles). Source localizations show the average regression weights obtained for the interaction effect in the significant time window, with a typical auditory profile. A post-hoc split between participants who did not recognize the foreperiod manipulation during debriefing (in yellow and those who did, in green) suggests that the cue-evoked difference in temporal predictability is larger in the second group. **B. Target-evoked activity** showed an interaction between foreperiod and predictability, with the strongest difference between predicted and non-predicted targets after short foreperiods, localized mainly to auditory areas. **C. Late target-evoked activity** differed between the pitch and duration discrimination tasks, showing stronger activity in auditory areas for pitch discrimination. The color code for the source activity reflects a relative scale, between the 95th (red) and 100th percentile (yellow).

**Table 1:**
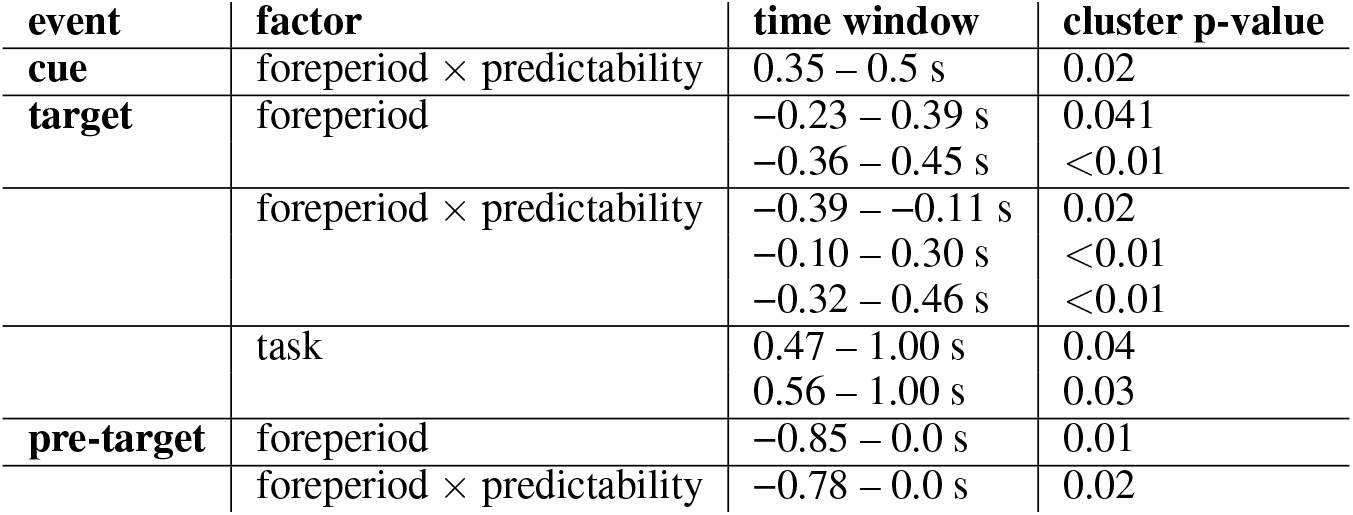
Statistical results, event related fields: Significant results from the group-level cluster permutation tests, computed for cue- and target evoked activity (magnetometers). Time window in which the respective contrast was significant and corresponding cluster p-value.

| event | factor | time window | cluster p-value |
| --- | --- | --- | --- |
| cue | foreperiod × predictability | 0.35 – 0.5 s | 0.02 |
| target | foreperiod | -0.23 – 0.39 s<br>-0.36 – 0.45 s | 0.041<br><0.01 |
| | foreperiod × predictability | -0.39 – -0.11 s<br>-0.10 – 0.30 s<br>-0.32 – 0.46 s | 0.02<br><0.01<br><0.01 |
|  | task | 0.47 – 1.00 s<br>0.56 – 1.00 s | 0.04<br>0.03 |
| pre-target | foreperiod | -0.85 – 0.0 s | 0.01 |
| | foreperiod × predictability | -0.78 – 0.0 s | 0.02 |

For the first-level regression, three factors were used: foreperiod, predictability, and task. We also tested for all two-way interactions, as well as the three-way interaction between foreperiod, predictability, and task. All predictors were zeroed. We computed the regression per time-point in the peri-stimulus time window (cue: 0.00 – 0.50 s, target: −0.50 – 1.00 s) and sensor. For the analysis of slow ramping activity, we removed the shortest foreperiod (0.5 s), and performed the statistical analyses on the interval from −1.0 – 0.0 s, time-locked to target onset.

On the group level, we computed t-tests, testing for each factor the regression weights obtained on the individual level against 0, with spatio-temporal cluster-based permutations for multiple comparison correction (Maris and Oostenveld, 2007). The threshold for inclusion in the cluster-test was p < 0.05 and the final acceptance threshold p < 0.05. Statistical results are displayed in Table 1, and Figures 3 and 4. We did not consider the factor foreperiod as a factor of interest, as main effects thereof trivially reflect the differences in stimulus timing. Below, will only discuss foreperiod effects in interaction with the factors predictability or task.

**Figure 4:**
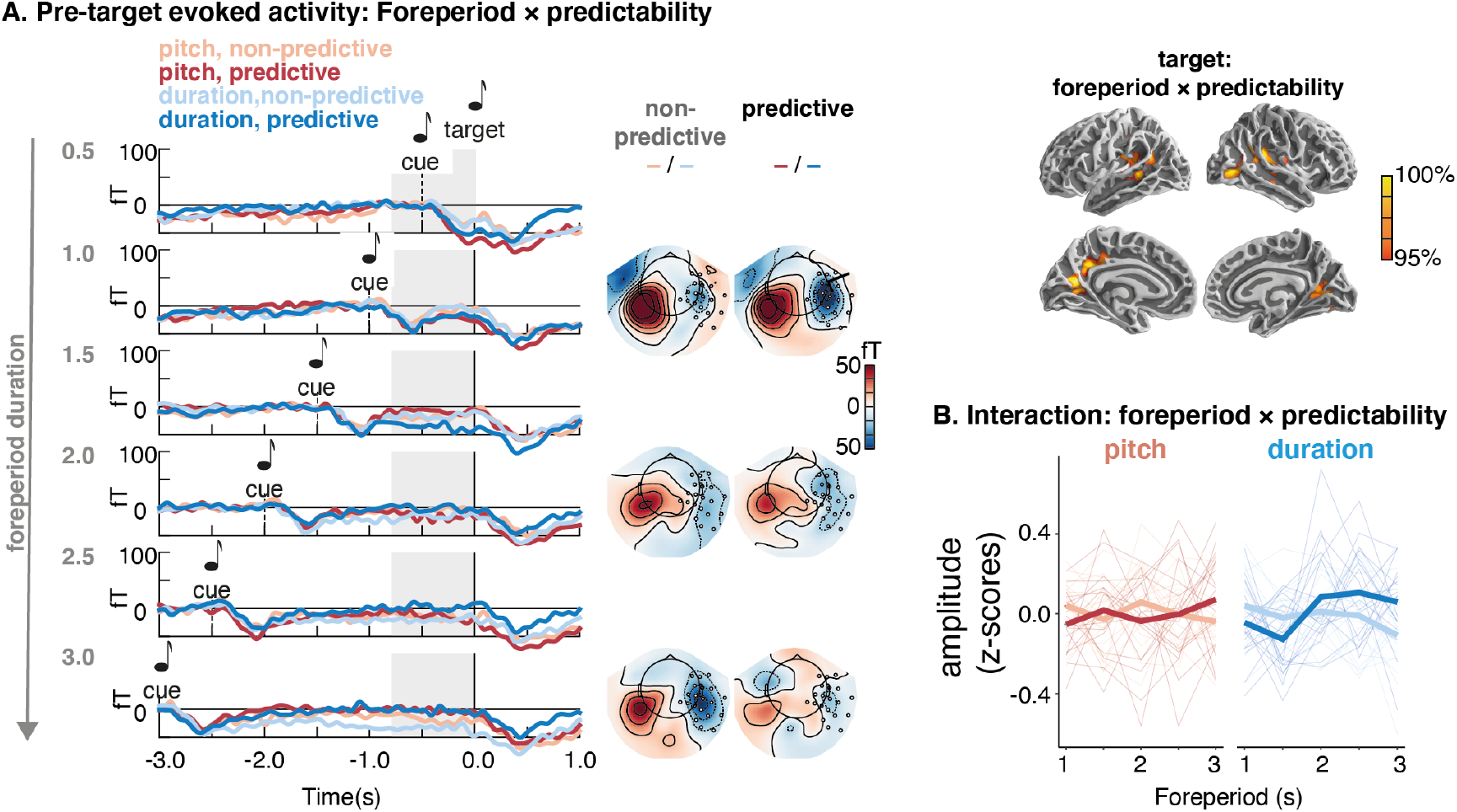
Slow deflections during the foreperiod: **A. Pre-target slow ramping activity**. Prior to target onset, a slow negative potential was observed, which reflects an interaction between foreperiod and predictability: at short foreperiods, activity was more negative for predictive versus non-predictive conditions, while at long foreperiods the relation flipped. **B. Interaction: Foreperiod** × **predictability:** The thin lines show single participants’ activity (z-scored), extracted from the significant sensors and time points in the group statistics. The thick lines depict the group average. The color code for the source activity reflects a relative scale, between the 95th (red) and 100th percentile (yellow).

#### Time-frequency analyses

First, we assessed general differences between the power spectra of the experimental conditions (presented in blocks), by computing Welsh’s power spectral density (PSD) over the 10 s long epochs, time-locked to cue onset, for frequencies from 0 – 40 Hz, with a frequency resolution of 0.1 Hz. PSD was computed on single epochs and then averaged per experimental condition (not shown as no differences were found). To test for differences in power spectra between the pitch and duration discrimination tasks, we computed paired t-tests per frequency and sensor on the difference between the two tasks. Cluster permutation tests were used to correct for multiple comparisons. These spectral analyses revealed no long-lasting differences in power between conditions.

Second, we assessed modulations of power during the foreperiod, focussing on the pre-target time window. To be able to include also the shortest foreperiods in this analysis, for which we found the strongest effects of temporal predictability in the behavioral data and evoked fields, we computed induced power by subtracting per condition (task, predictability, and foreperiod) the trial-average time domain response from each trial’s data. To avoid edge effects, we appended the left side of the epochs (−3.5 – 0.0 s with their flipped version: 0.0 – −3.5 s). Then we computed time-frequency transformations, using multitapers (bandwith = 2.0) on 40 linearly spaced frequencies from 0.5 – 40.0 Hz in steps of 1 Hz, with varying cycles per frequency (2 – 10, linearly spaced). Power estimates were baseline corrected by manually z-scoring each epoch with respect to its pre-cue baseline (−0.5 – −0.2 s before cue onset). The manual correction was necessary due to the varying foreperiods.

As for the time-domain activity, statistics were computed on two levels. Per participant, we computed, per time-frequency point and sensor, a linear regression predicting per-trial power, with the factors foreperiod, predictability, and task, as well as their possible interactions. A three-dimensional cluster test per regressor over frequency, time points, and sensors was computed in FieldTrip (Matlab 2020a), using similar parameters as for the time domain data. Testing was restricted to the pre-target time window (−0.5 – 0.0 s). Statistical results are displayed in Table 2, and Figure 5.

**Figure 5:**
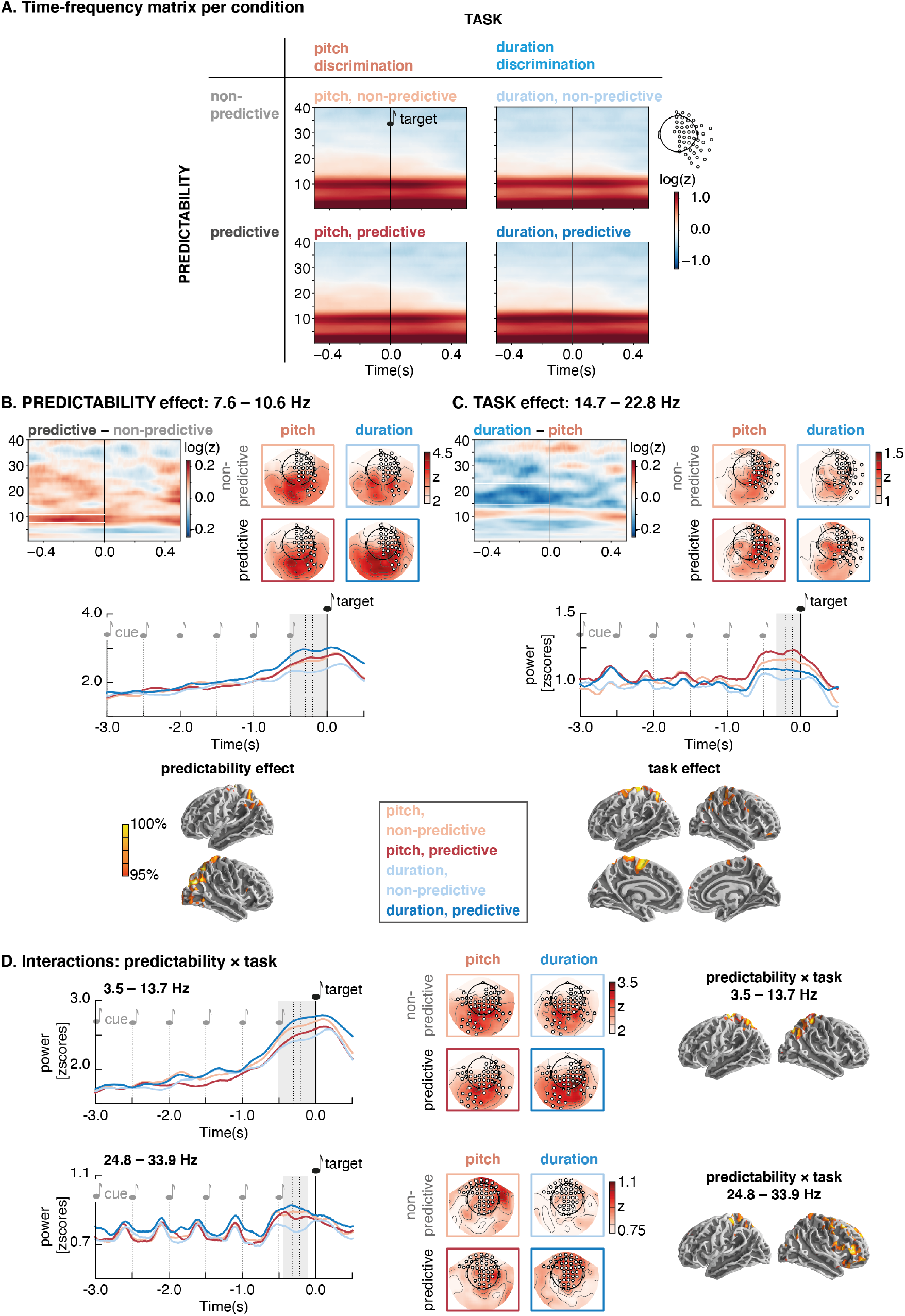
**A. Power per time point and frequency** for all four conditions (baseline corrected), extracted at the joint group of sensors depicted in B and C. **B. Main effect of predictability:** Temporal predictability was indexed by a stronger increase in alpha power prior to target onset, with sources in parietal areas, reaching into visual cortices. The top left panel shows the time-frequency matrix of the difference between the predictive and non-predictive conditions. The topographies show the distribution of alpha power across sensors per condition, with stronger alpha power in the predictive conditions of both tasks. The middle panel shows the alpha power time courses for all four conditions, time-locked to target onset (the gray shade marks the statistically significant time window), averaged over all sensors included in the significant cluster. Source plots show average regression weights for the task effect in the alpha band, localized to parietal and visual areas. **C. Main effect of task:** Induced power in the beta range showed a task difference, namely it was higher in the pre-target time window for the pitch discrimination compared to the duration discrimination task. The top panel shows the time-frequency matrix of the contrast between duration discrimination and pitch discrimination. The topographies show the distribution of beta power across sensors per condition, with stronger beta activity over right central sensors in the pitch discrimination task. The middle panel shows the beta power time course for all four conditions, time-locked to target onset (the gray shade marks the statistically significant time window), averaged over all sensors included in the significant cluster. Source plots show average regression weights for the task effect in the beta band, localized to motor and parietal areas. **D. Interactions between predictability and task:** Top row: interaction between predictability and task in the theta/alpha range with higher power for the conditions pitch non-predictive, and duration predictive. The interaction was found over parietal areas, comparable to those shown for the predictability effect in panel B. Bottom row: interaction between predictability and task in the higher beta range, with the lowest power in the non-predictive blocks of the pitch discrimination task, over central and frontal sensors. The color code for the source activity reflects relative activity between the 95th (red) and 100th percentile (yellow).

**Table 2:** Statistical results, induced power: Significant results from the group-level cluster permutation tests, computed in the foreperiod window (pre-target). Time window in which the respective contrast was significant, frequencies, and corresponding cluster p-value.

| event | factor | time window | frequencies | cluster p-value |
| --- | --- | --- | --- | --- |
| target | foreperiod | -0.5 – 0.00 s | 0.5 – 18.7 Hz | <0.01 |
| target | task | -0.32 – 0.00 s | 14.7 – 22.8 Hz | 0.04 |
|  | predictability | -0.50 – 0.00 s | 7.6 – 10.6 Hz | 0.03 |
| | task × predictability | -0.50 – 0.00 s | 3.5 – 13.7 Hz | <0.01 |
| | task × predictability | -0.43 – 0.12 s | 24.8 – 33.9 Hz | <0.01 |

#### Source Reconstruction

Individual head models were computed using the anatomical T1-weighted images recorded for each participant. The volumetric segmentation of participants’ anatomical MRI and the cortical surface reconstruction was performed with the FreeSurfer software (Fischl, 2012). Single layer individual head models were computed using the boundary-element method (BEM), using each participant’s anatomical MRI. The coregistration of the MEG data with the individual’s structural MRI was carried out by manually realigning the digitized fiducial points with MRI slices (using MNE python’s command line tools). Second, an iterative refinement procedure was used to realign all digitized points with the individuals’ scalp.

The noise covariance matrices were estimated using the empty room recordings for each participant, after applying the same low-pass filter as to the task data. Source reconstruction illustrating oscillatory power effects used noise covariance matrices bandpass-filtered to the frequency band for which a significant cluster was found.

We computed individual forward solutions (surface models; spacing = ico 5), resulting in 10,242 voxels per hemisphere. We then projected per participant the first-level regression weights from the sensor space analyses described above onto the surface, using dynamical Statistical Parametric Mapping (dSPM, Dale et al., 2000), with partially fixed dipole orientation (0.2), and depth weighting of 0.8. Individual surface activation maps were aligned to FreesSurfer’s ‘fsaverage’ template brain, and averaged over all participants.

## Results

### Behavioural responses

#### Temporal predictability improves the discrimination of pitch, but not of duration

Participants’ pitch discrimination accuracy was enhanced in the temporally predictive condition compared to the non-predictive condition (non-predictive: 0.71 (SD = 0.11), predictive: 0.73 (SD = 0.13), T(22) = −2.88, p < 0.01, Bayes Factor (BF) = 7.13). Duration discrimination accuracy did not differ significantly between predictive and non-predictive conditions (0.78 (SD = 0.09) vs. 0.77 (SD = 0.12), T(22) = −0.66, p = 0.51, BF = 0.35), see also Figure 2A. We observed a significant difference in accuracy between the pitch and duration judgement tasks, albeit with an inconclusive Bayes Factor (pitch: 0.72 (SD = 0.12) duration: 0.78 (SD = 0.09), T(22)= −2.18, p = 0.04, BF = 1.91).

#### Temporal predictability speeds up response times

Response times were faster in the temporally predictive condition in both tasks (pitch discrimination: 0.56 (SD = 0.12) vs. 0.52 s (SD = 0.11), T(22) = 4.80, p < 0.001, Bayes Factor (BF) = 941.67; duration discrimination: 0.54 (SD = 0.11) vs. 0.51 s (SD = 0.09), T(22) = 3.60, p < 0.01, Bayes Factor (BF) = 37.01; see Figure 2B). We also found a significant interaction between temporal predictability and foreperiod in both tasks (pitch: F(5,22) = 4.71, p < 0.001; duration: F(5,22) = 2.34, p = 0.046), indicating that the difference in response times due to temporal predictability occurred predominantly at the short foreperiods. Post-hoc tests showed that in the pitch-discrimination task, the difference was significant at foreperiods 0.5 and 1 s (p < 0.001, p < 0.01; all other p > 0.25), and in the duration discrimination task at foreperiods 0.5, 1, and 1.5 s (p = 0.02, p < 0.001, p < 0.001, all other p > 0.35).

#### Explicit timing behavior

To verify that participants had engaged in explicit timing, even in the predictive blocks when the same interval was presented on each trial, we assessed duration discrimination responses over trials (Figure 2C). As expected, the average response (‘short’ = −1, ‘long’ = 1) gradually increased with foreperiod, in the non-predictive and predictive blocks (Figure 2C, top), indicating that duration judgements reflected the presented durations. Also, individual responses (depicted by the thin lines in the top panel) showed similar variability in predictive and non-predictive blocks, ruling out the alternative assumption that participants repetitively pressed the same button in the predictive blocks, without timing each interval. Similarly, the standard deviations (computed per participant over trials) was highest in for the intermediate foreperiods. Paired t-tests between the standard deviations of the non-predictive and predictive conditions per foreperiod revealed (marginally) larger standard deviations for the non-predictive blocks, for the foreperiods 0.5, 1.0, and 1.5 s (0.5 s: T(22) = 2.4, p = 0.02; 1.0 s: T(22) = 1.8, p = 0.09; 1.5 s: T(22) = 2.8, p = 0.01; all other p > 0.16). This result indicates that participants might have benefited from the predictability when judging the duration of the shorter foreperiods.

Additionally, we tested the responses in the explicit timing task for linear trends over trials, which would be indicative of a shift in the internal reference throughout the block (fits shown in Figure2C, bottom right). T-tests against 0 revealed no significant linear fits in the non-predictive condition (all p > 0.2), and a significant negative slope in for the 2 s-foreperiods in the predictive condition (T(22) = −2.7, p = 0.01, as well as a marginal effect for the 2.5 s foreperiod T(22) = −2.1, p = 0.06; all other p > 0.14). These findings indicate that a reference shift might have occurred in the predictive blocks with ambiguous durations, providing further evidence that participants did engage in explicit timing throughout these blocks.

#### Temporal predictability improves pitch discrimination sensitivity

In line with our main hypothesis, we addressed whether behavioral sensitivity, indexed by the slope of the psychometric functions (shown in Figure 2D) did improve with temporal predictability. The slopes of the psychometric functions were significantly steeper in the pitch discrimination task for the temporally predictive condition as compared to the non-predictive one (T(22) = −2.90, p < 0.01, BF = 7.49). This was not observed in the duration discrimination task (T(22) = 0.05, p = 0.96, BF = 0.30), see Figure 2E. We observed no effects on the thresholds of the psychometric functions.

The slope and accuracy effects were significant in the group of 19 participants for whom we analysed the MEG data (accuracy: p = 0.01, BF = 5.96; slope: p = 0.02, BF = 3.75).

In sum, temporal predictability improved the performance in the pitch discrimination task, as indexed by accuracy and behavioral sensitivity (slope), but did not affect performance in the duration discrimination task. The absence of a performance increase in the duration discrimination task appeared surprising at a first glance – we had expected to see a facilitation of explicit timing due to the repetition of foreperiods over blocks. It is important to mention here that no feedback was given, meaning that participants could not correct their temporal judgements with respect to an objective duration. Furthermore, no objective reference interval was presented per trial, with the intention to keep the stimulation in the pitch and duration discrimination tasks the same. Hence, participants relied on their subjective estimates of duration. Furthermore, we think that the absence of performance differences between the non-predictive and predictive blocks in explicit timing confirms that temporal predictability was processed mainly implicitly and therefore did not affect explicit duration discrimination.

#### No correlations between implicit and explicit timing performance

We found no significant correlations between the measures of implicit and explicit timing. Correlating the ratio between predictive and non-predictive conditions for slope, accuracy, and response time in the pitch discrimination task with the respective ratios in the duration discrimination task yielded no significant outcomes (slopes: r = −0.32, p = 0.13, BF = 0.50; RT: r = 0.35, p = 0.10, BF = 0.52; accuracy: r = 0.04, p = 0.87, BF = 0.52).

#### Post-experimental questionnaire

Participants reported no significant differences in the difficulty of the pitch and the duration discrimination tasks, as assessed in the post-experimental questionnaire (mean 5.50 vs. 4.43 on a scale of 1 – 10; SD: 2.16 vs. 2.12; T(19) = 1.7, p = 0.10, BF = 0.97). Three participants did not, or only partially respond to the questionnaire.

All participants noticed that the time intervals varied, but only one spontaneously described that there were identical series of foreperiods. After being told that the intervals were fixed during some blocks, thirteen participants confirmed they had noticed this, seven of which only noticed it in the duration discrimination task, but not in the pitch discrimination task. Six participants claimed that fixed foreperiods had helped them in the pitch discrimination task, and two explicitly stated that it helped most at the shortest foreperiods. Thirteen participants reported that fixed durations helped them in the duration discrimination task. In sum, participants were not able to spontaneously describe the foreperiod manipulation, which shows that they were not fully aware of the temporal predictability. However, the fact that a substantial number of participants confirmed noticing the fixed foreperiod blocks after it was described to them indicates that they may have gained partial awareness of the predictability manipulation.

### Modulations of auditory activity by temporal predictability and task

#### Cue-evoked activity reflects temporal predictions

At latencies following the cue, we observed an interaction between predictability and foreperiod: the auditory response to temporally predictive cues was stronger than to non-predictive cues, and this effect was strongest at the shortest foreperiods (0.35 – 0.50 s, p = 0.02, Figure 3A, Table 1). Post-hoc tests revealed that the interaction was significant for the shortest foreperiod (0.5 s, T(18) =-4.27, p < 0.001, BF = 72.56, all other p > 0.24). For the 2 s foreperiod, the effect was inversed (T(18) =-2.75, p < 0.013, BF = 4.16, no correction for multiple comparisons).

Since the relatively late latency of the effect could indicate that temporal predictions conveyed by the cue were processed consciously (Sergent et al., 2005), we performed a post-hoc split of participants in two groups with respect to whether they had claimed to recognize the foreperiod manipulation or not (Figure 3A, inset bottom right). Although the small number of participants did not allow for a robust statistical comparison, the effect appeared to be larger in the group of participants who noticed the predictability manipulation (in green, larger average absolute regression weight; confidence interval does not include 0).

#### Target-evoked activity is affected by temporal predictability and task

In the target time window, we observed an interaction between foreperiod and predictability in three bilateral significant clusters ranging from −0.40 to 0.46 s post-target onset (p = 0.03, Table 1 and Figure 3B). The effects of foreperiod and of predictability started before target onset, thus likely overlapping with the cue-evoked differences.

The interaction between foreperiod and predictability consisted in larger evoked activity following predictive foreperiods as compared to non-predictive ones: the shorter the foreperiod, the more marked this difference was, appearing particularly salient on the topographic maps (Table 1, Figure 3B). Post-hoc tests revealed that the interaction was driven by the shortest foreperiod (0.5 s, T(18) =-4.79, p < 0.001, BF = 200.36, all other p > 0.12). Again, we observed a reversed effect (non-predictive > predictive) at the 3 s foreperiod (T(18) =-2.84, p = 0.01, BF = 4.85).

The source localization of this effect showed, as for the cue-evoked effect, the implication of early auditory areas. The interaction between foreperiod and predictability aligns well with the similar interaction found in the response times. Altogether, these observations suggest that temporal predictability was extracted regardless of the task, and had the strongest effect on the processing of early target tones.

Furthermore, following the target we observed a late main effect of task (two clusters, 0.47 – 1.00 s, p = 0.04, Figure 3C), which resulted in a more sustained response in auditory areas for the pitch discrimination task as compared to the duration discrimination task.

In an additional analysis devised to assess slow ramping activity prior to target onset, we observed an interaction between foreperiod and predictability (−0.78 – 0.0 s, p = 0.02, Table 1, see Figure 4): during the foreperiod, there was a sustained negative deflection, which was stronger in the predictive compared to non-predictive conditions at short foreperiods, and less negative at the longer foreperiods. The shape of this activity resembles the contingent negative variation (CNV/ CMV), but the generators were in auditory areas.

#### Modulation of induced alpha and beta power during the foreperiod

In the time-frequency analyses, we focused on the pre-target time window, to assess the effects of temporal predictability and of task, when participants were either *implicitly timing* in anticipation of the upcoming target in the pitch task, or *explicitly timing* in anticipation of the tone marking the end of time interval in the duration task. As both, behavioral data (RT) and evoked fields showed the strongest benefit of temporal predictability for short foreperiods, we used all foreperiods in this analysis and focused on induced power, to avoid confounding cue-evoked effects with pre-target modulations. Statistical results are reported in Table 2.

Temporal predictability was found to modulate pre-target alpha power (7.6 – 10.6 Hz, −0.50 – 0.00 s pre-target, p = 0.03; see Figure 5B), with higher alpha power in temporally predictive conditions over the parietal and visual cortices. The parietal regions also showed an interaction between task and predictability, in a broader frequency range (3.5 – 13.7 Hz, −0.50 – 0.00 s pre-target, p < 0.01; Figure 5D, upper row). Post-hoc paired t-tests on the average amplitudes extracted from the significant cluster (time points, sensors) showed that the interaction resulted from a reversal of the direction of the predictability effect in the pitch discrimination task (T(18) = 2.87), p = 0.01, BF = 5.12) compared to the duration discrimination task (T(18)=-4.41), p < 0.001, BF = 96.35). The task difference was significant in the non-predictive blocks (pitch versus duration: T(18) = 3.26, p < 0.01, BF = 10.51), but not in the predictive blocks (T(18)=-1.59, p = 0.13, BF = 0.69).

A main effect of task was observed in the beta frequency range (14.7 – 22.8 Hz, −0.32 – 0.00 s pre-target, p = 0.04, see Figure 5C), indicating that the pitch task elicited higher beta power during the foreperiod than the duration task. The statistical significance occurred over the right central sensors, and source reconstruction of the regression weights revealed sources mainly in central (motor) areas, including the left SMA, and parietal areas, bilaterally.

As we had observed a significant task difference in behavioral accuracy, we assessed whether the main effect of task in the beta band was driven by task difficulty. The correlation between between the task difference in accuracy and the task difference in beta power was marginally significant (*ρ* = 0.43, p = 0.06), but driven by four participants who had a large difference in task difficulty, and no or a positive difference in beta power (compared to the negative difference observed in the main effect). After removing those four participants the correlation was not significant (*ρ* = −0.26, p = 0.33). This confirms that the task difference in the beta band was not driven by differences in task demands.

A second interaction occurred in the upper beta band (24.8 – 33.9 Hz, −0.43 – −0.12 s pre-target, p < 0.01; Figure 5D, bottom row). Post-hoc tests showed that the predictive and non-predictive conditions differed significantly only in the duration task (pitch: T(18) = 0.64, p = 0.53, BF = 0.29; duration: T(18) = −4.89, p < 0.01, BF = 239.88). The task difference was significant but reversed in direction in the non-predictive blocks (pitch versus duration: T(18) = 3.81, p < 0.01, BF = 29.76), as well as in the predictive blocks (T(18) = −2.69, p = 0.02, BF = 3.70). The sources were more wide-spread over central and frontal areas, and the frequency band did not overlap with the main effect of task.

None of the above effects showed a significant interaction with the foreperiod (predictability × foreperiod: p = 0.15 (22.8 – 27.8 Hz); task × foreperiod: p = 0.10 (23.8 – 27.8 Hz); 3-way interaction: p = 0.06 (3.5 – 8.6 Hz)). Exploratory analyses were carried out to test for differences in inter-trial phase coherence between conditions, as well as differences in phase-amplitude coupling strength, but yielded no consistent results.

## Discussion

In this study, we asked whether implicit temporal prediction and explicit duration judgement rely on shared versus separate timing mechanisms. We induced implicit timing in an auditory foreperiod paradigm by manipulating the foreperiod to be either temporally predictive (constant) or not (variable). Participants received two consecutive task instructions: pitch discrimination (indirect measure of implicit timing) and duration discrimination (direct measure of explicit timing), while their brain activity was recorded with magnetoencephalography (MEG). Participants were not informed about this manipulation, and post-experimental debriefing showed that they were not explicitly aware of it. To engage participants in explicit timing, we asked them to perform a duration discrimination task on the foreperiod intervals.

The results show that participants systematically formed temporal predictions in both tasks, speeding up response times and enhancing the auditory evoked response. However, behaviourally, only pitch discrimination performance benefited from the temporal predictability. Attentional orienting in time during predictive foreperiods was indexed by an increase in alpha power over parietal and visual areas. Pre-target induced beta power in sensorimotor and parietal areas was increased during pitch discrimination compared to duration discrimination, suggesting that beta power here reflects implicit timing. Interestingly, we observed no distinct neural signatures when participants were asked to provide explicit duration judgements (duration discrimination), compared to implicit timing alone (pitch discrimination).

### Temporal predictions are formed irrespective of the task, but only affect pitch discrimination performance

The behavioral findings (Figure 2) clearly show that our manipulation of the foreperiod induced implicit timing. Slower response times at short foreperiods are commonly reported in variable foreperiod designs (Cravo et al., 2011; Niemi and Näätänen, 1981; Nobre and van Ede, 2018), indicating that participants automatically extract the temporal statistics of the inputs (here: the range of foreperiods; Droit-Volet and Coull, 2016; Martin et al., 2017). A novel finding we report here is that when combining variable and fixed foreperiods, the temporal predictability allowed to overcome the slowing of responses at short foreperiods.

While response times reflected the extraction of temporal predictability irrespective of the task, pitch discrimination sensitivity increased after temporally predictive foreperiods, replicating a previously published finding (Herbst and Obleser, 2019). Duration discrimination performance did however not benefit from temporal predictability. Interestingly, the response time effects, and MEG results show that temporal predictability was extracted, also in the duration judgement blocks, but apparently the implicit knowledge of temporal statistics did not improve the explicit duration judgements. This finding points to a difference between implicit temporal predictions that serve to orient attention in time, and explicitly encoded durations.

Importantly, the duration judgements showed that participants did engage in explicit timing, even when the same foreperiod interval was presented repeatedly in the predictive blocks. Duration judgements showed comparable patterns of mean and variability in the non-predictive and predictive blocks (Figure 2C). Furthermore, we observed a linear trend in the responses in the predictive block with 2 s foreperiods indicating that participants were more likely to judge the interval as ‘long’ in the beginning, and as ‘short’ in the end of a block. This shift suggests an adjustment of the internal reference used for the comparison, an established effect in explicit timing (Dyjas et al., 2012). Here, during predictive blocks, the moving average of previously presented durations might shift in the direction of the foreperiod used on that block, hence increase in the case of the 2 s or 2.5 s durations. Thus, the comparison foreperiod would appear shorter, especially towards the end of the block. This effect is more likely to affect the ambiguous intervals, than the ones for which the discrimination is clear. Taken together, the observed patterns of variability and the drift for the intermediate intervals support the assumption that participants encoded and judged the foreperiod interval on each trial even in the predictive blocks.

Notably, we observed no correlations between implicit and explicit timing performance, in line with a majority of previous studies that reported dissociable behavioral patterns when comparing implicit and explicit timing across individuals (Droit-Volet and Coull 2016; Droit-Volet et al. 2019; Los and Horoufchin 2011; Mioni et al. 2018; but see Coull et al. 2013, long intervals). In line with the absence of an improvement of explicit timing by predictability, this finding suggests that the implicit temporal predictions are not directly translated to duration estimates.

Overall, the behavioral results show that the human brain is tuned to efficiently extract implicit temporal statistics of sensory environments, which here, were present in both tasks due to the balanced design. Yet, the behavioral results indicate that temporal predictability is only used implicitly, to benefit sensory processing and reduce response times, but does not affect the explicit duration judgements.

### Implicit timing enhances the auditory response to predictive and predicted stimuli

The enhanced auditory response to predicti*ve* cues and predict*ed* targets (Figure 3) confirms that temporal predictability anticipates and enhances the auditory representation of sound (Jaramillo and Zador, 2011; Miniussi et al., 1999), regardless of whether the sound’s properties are directly task relevant (as in the pitch discrimination task), or whether the sound only serves as a temporal marker (as in the duration discrimination task). Furthermore, the predictability effect occurred in interaction with the foreperiod (driven by the shortest foreperiod), which confirms that temporal predictions are conditionally updated over time (Giersch et al., 2016; Herbst et al., 2018; Nobre et al., 2007), and therefore more useful to anticipate early target onsets. In light of the recent debates on the effects of attention versus prediction (de Lange et al., 2018; Heilbron and Chait, 2018; Press et al., 2020; Todorovic et al., 2015), our results align with an attentional enhancement of auditory processing: here temporal predictions allowed to orient attention in time, to the most likely moment of target occurrence.

We also observed anticipatory auditory activity to be modulated by temporal predictability in interaction with the foreperiod (Figure 4). Such anticipatory auditory activity was previously reported during explicit timing of auditory stimuli (Bendixen et al., 2005; Picton et al., 1978; van Wassenhove and Lecoutre, 2015), and when auditory stimuli contain implicit temporal regularities (Sohoglu and Chait, 2016), and is in line with sensory timing models (Bueti, 2011; Eagleman and Pariyadath, 2009), which hold that the neural populations that process the sensory stimuli also contribute to representing its temporal features.

### Induced alpha and beta power carry distinct signatures of attentional orienting and implicit timing

#### Alpha power as a signature of attentional orienting

Distinct signatures of orienting attention in time were observed in parietal/ visual alpha power increasing prior to the predicted compared to the non-predicted targets in both tasks (Figure 5B). The modulation of alpha power prior to temporally predicted stimuli is a classical finding, usually surfacing as a decrease when the stimuli are visual (Ede et al., 2016; Rohenkohl and Nobre, 2011; Solís-Vivanco et al., 2018; van Diepen et al., 2015). In the auditory domain, attentional modulations of alpha power often show the reversed direction, namely an increase in alpha over visual/parietal areas (Schneider et al., 2021; Strauß et al., 2014), interpreted as a suppression of task-irrelevant (e.g. visual) processes (Jensen and Mazaheri, 2010; Klimesch et al., 2007). The stronger increase in alpha power during predictive foreperiods aligns nicely with those earlier findings, and replicates a previous finding of increased alpha power prior to temporally predictable auditory stimuli (Herbst and Obleser, 2017). The parietal sources of the alpha power effects are in line with a previously reported involvement of these areas in temporal prediction (Coull and Nobre, 2008; Meindertsma et al., 2018; Visalli et al., 2019).

We also observed an interaction between predictability and task in at least partially overlapping anatomical regions compared to the main effect of predictability, but in a broader frequency band, from low theta to high alpha (Figure 5D), likely reflecting an on-top modulation of the predictability effect by task. The parietal sources of the interaction (compared to the parieto-visual effect of predictability) suggests the involvement of the dorsal attentional orienting network (Corbetta and Shulman, 2002; Petersen and Posner, 2012), also supported by the broader frequency range found to be modulated (Cavanagh and Frank, 2014; Fox et al., 2006; Haegens et al., 2011; Palva and Palva, 2011; Sadaghiani et al., 2010). We speculate that this gradual decrease in alpha power over conditions indicates differential allocation of attentional resources to elapsed time throughout the foreperiod.

#### Beta power as a signature of implicit timing

When comparing the two timing tasks, we did observe a difference in pre-target induced beta power over sensorimotor and parietal areas, including the SMA (Figure 5C): beta power was increased during pitch discrimination compared to duration discrimination. Lower power in the explicit timing task might appear counter intuitive at first, as previous studies found beta-power to increase when comparing an explicit timing task to non-timing task (Kulashekhar et al., 2016; Spitzer et al., 2014). However, these studies did not modulate implicit timing, and they used visual and tactile comparison tasks. The study by Kulashekhar et al. reports a relative suppression of beta power compared to baseline, both in the visual and the timing task, but the suppression is stronger in the non-timing task, resulting in a positive difference. Furthermore, in explicit non-rhythmic motor timing, a transient increase in beta power after the self-initiated onset button press has been found to reflect the duration the participant will produce (Kononowicz and Rijn, 2015; Kononowicz et al., 2018), thought to index the inhibition of a state-dependent timing network. In our results, the modulation of beta power occurred throughout the foreperiod, with no distinct temporal profile, suggesting that, without a clear motor component in the explicit timing task, implicit timing engages oscillatory dynamics in the beta range more so than explicit timing.

Increased beta power during implicit timing was previously reported during rhythmic temporal prediction (Fujioka et al., 2012; Morillon and Baillet, 2017), and predictive foreperiods (Meindertsma et al., 2018), and aligns more generally with the involvement of beta oscillations in top-down predictions (Arnal, 2012; Bastos et al., 2015; Engel and Fries, 2010; Spitzer and Haegens, 2017). In our case, beta power was increased in both conditions of the pitch discrimination task, where temporal predictions appeared beneficial for task performance. In line with the behavioral findings, this confirms that in the variable foreperiod conditions, participants conditionally updated their temporal predictions with elapsed time (Bueti et al., 2010; Giersch et al., 2016; Herbst et al., 2018; Janssen and Shadlen, 2005).

### Shared or separate timing mechanisms?

Taken together, our results suggest that the brain flexibly uses overlapping time coding mechanisms in implicit and explicit timing situations, depending on the temporal statistics of the inputs and the task requirements (see also van Ede et al., 2020), rather than deploying specialized modules for either timing task.

A noteworthy outcome of this study is the absence of distinct brain dynamics emerging during the to-be-timed interval when attention is directed *to* time during explicit timing (duration discrimination), compared to implicit timing (pitch discrimination), in other words, the absence of a positive task difference. This divergence from the previous fMRI work (Coull et al., 2013) is likely due to the different neural dynamics targeted by fMRI versus MEG, and the focus on a fine-grained temporal resolution in our study. Tentatively, the absence of distinct neural dynamics during explicit timing in this study could indicate that elapsed time is encoded by a common mechanism during implicit and explicit timing, and subjected to separate read-out mechanisms per task. Such task-independent encoding of time has been found in several brain areas, for instance a recent report showed precise encoding of elapsed time in rats’ dorsal striatum, which was, however, unrelated to subjective duration perception (Toso et al., 2021). This suggests that downstream readers are needed to transform the signals which encode elapsed time to task-relevant information (Buzsáki, 2010; van Wassenhove, 2009). Possibly, for the implicit timing task, the relevant readout consists in a prediction error (Meirhaeghe et al., 2021), while for the purpose of explicit timing a duration is required (Kononowicz et al., 2018; van Wassenhove, 2009). Different read-out mechanisms would also allow to account for the observed differences in behavior between the two timing tasks. One reason for not observing any neural signatures related to the readout in this study might be that this process is not necessarily locked to a specific time point (target onset).

### Comparing implicit and explicit timing: limitations

Comparing an implicit cognitive process, such as implicit timing, with an explicit one is a challenging endeavour, as it requires another sensory task during which the implicit process takes place (here: the pitch discrimination task). While we are aware that the choice of that sensory task might have affected the results, several measures were taken to ensure the comparability between conditions: notably, we used empty intervals delimited by two sensory events to keep the stimulation exactly the same in both tasks and avoid any sensory processing during these intervals on which we focussed our comparisons of neural dynamics. It should also be noted that performance was significantly better in the duration discrimination task compared to the pitch discrimination task. However, this difference did not correlate significantly with the observed difference between tasks in beta power.

To achieve a balanced design, the study had a 2 × 2 manipulation, with the factors predictability and task. The main effects found for both factors suggest an implementation by separate neural dynamics (alpha, and beta oscillations), but the interactions also show differences between the individual conditions. Necessarily, our design contains a sub-selection of timing tasks, and future studies should vary tasks and sensory modalities to test the generality of those results.

Notably, the non-predictive condition in the pitch discrimination task was intended to reflect a temporally neutral condition (Coull et al., 2013), with no incentives to engage in timing (Figure 1B). However our results clearly show that the brain constantly extracts temporal statistics from sensory environments (Jazayeri and Shadlen, 2010; Nobre et al., 2007): the interaction effect between predictability and foreperiod indicate that the temporal predictability mainly benefited sensory processing and motor preparation at the short foreperiods, and that participants updated their predictions with elapsed time, even in the non-predictive blocks.

### Conclusions

In this study, temporal predictions systematically enhanced auditory processing, but did not translate to explicit duration judgements. In line with previous studies, alpha power indexed the orienting of attention to temporally predicted events. Increased beta power surfaced as a signature of implicit timing in the pitch discrimination task to which temporal predictions were also relevant for performance. However, we did not observe distinct neural dynamics during the foreperiod when attention explicitly directed to time. Our work thus indicates that implicit timing shapes the behavioral and sensory response in an automatic way, and is reflected in oscillatory neural dynamics, while the translation of implicit temporal statistics to explicit durations remains inconclusive, possibly due to the more abstract nature of the task.

## Conflict of interest statement

The authors declare no conflict of interest.

## Acknowledgements

The authors would like to thank the members of the Cognition and Brain Dynamics Team and Jens Kreitewolf for helpful comments on the implementation of the study, and the members of UNIACT at NeuroSpin for help with participant recruitment and preparation, as well as Leila Azizi and Pooja Prabhu for help during the recordings. The study was funded by a DFG Grant HE 7520-1/1 to SKH, and ANR-16-CE37-0004-04 AutoTime to VvW.

## Appendix

### Debriefing questionnaire

How tiring did you find the experiment on a scale from = not tiring at all, to 10 = very tiring?
Which aspects were tiring?
How difficult was it for you to separate the tones from the background noise on a scale from 1 = not difficult at all, to 10 = very difficult?
How difficult was it for you to judge the pitch on a scale from 1 = not difficult at all, to 10 = very difficult? How difficult was it for you to judge the duration on a scale from 1 = not difficult at all, to 10 = very difficult? Did you realize that the time interval between the two tones was variable? If yes, can you describe the variability?
In fact, there were blocks were the interval was always the same, and others were it varied. Did you notice that? Did you think this helped you with performing the tasks (answer separately for pitch and duration)? Do you have other remarks about the experiment?
Do you play a musical instrument? If yes, which, and for how many years?

## Footnotes

1 https://github.com/brainthemind/CogBrainDyn_MEG_Pipeline

## Notes

### Competing Interest Statement

The authors have declared no competing interest.

